# Generation of Synthetic scRNA-seq-Like Transcriptomes Using a Generative Adversarial Network from RNA-seq Data

**DOI:** 10.1101/2025.10.09.681449

**Authors:** Sujit Silas Armstrong Suthahar, Dan Ruan

**Affiliations:** University of California, Los Angeles, Department of Bioengineering, Los Angeles, 90095, USA

## Abstract

Next-generation sequencing (NGS) technologies have become integral for high-throughput transcriptomic studies. Among these, single-cell RNA sequencing (scRNA-seq) is especially valuable for quantifying gene expression at the individual cell level, enabling the identification of rare cell populations and cellular differentiation pathways. However, the high cost of scRNA-seq often limits its broader application. Bulk RNA sequencing (RNA-seq) provides a more affordable alternative but lacks the single-cell resolution needed to elucidate cellular heterogeneity. Here, we present a cycle-consistent generative adversarial network (cycleGAN) approach to generate synthetic single-cell-like transcriptomes from bulk RNA-seq data. By adversarially training two sets of generators and discriminators, our framework attempts to learn the relationship between bulk and single-cell transcriptome distributions. Although this approach does not replace real scRNA-seq experiments, it can be a usefult tool to generate synthetic single-cell-like data for preliminary exploratory investigations and other machine learning applications. We further discuss the performance, limitations, and ethical considerations of our method.

## Introduction

Single-cell RNA sequencing (scRNA-seq) provides powerful insights into cellular heterogeneity, enabling researchers to identify rare cell types, study cell-cell interactions, and unravel complex gene expression patterns in tissue and blood samples^1,2^. However, the high cost of scRNA-seq—estimated at roughly $21,300 for profiling 100,000 cells, including library preparation^3^— can be a limiting factor for many laboratories.

Bulk RNA sequencing (RNA-seq) offers a more affordable approach to transcriptomic analysis, yet it does not capture gene expression at the single-cell level. To bridge this gap, we propose a Cycle-Consistent Generative Adversarial Network (cycleGAN) that learns to transform bulk RNA-seq data into synthetic scRNA-seq-like profiles. While it may not be a substitute for experimentally generated scRNA-seq, it serves as a cost-effective tool for exploratory analyses, hypothesis generation, and training of downstream machine learning and deep learning models.

Although there are several algorithms for generating synthetic scRNA-seq data, such as scDesign^4–6^, Splatter^7^, and ZINBWaVE^12^, these methods typically rely on statistical models and require existing scRNA-seq dataset from which they generate synthetic profiles. Our cycleGAN-based approach attempts a direct translation from bulk RNA-seq to synthetic single-cell profiles. The fundamental assumption of our model is that some aspects of the underlying single-cell distribution can be inferred from bulk data when representative single-cell data for training are available. However, we stress that bulk RNA-seq does not generally contain the full resolution of true single-cell transcriptomes, and our method does not guarantee recovering new or unseen cellular subpopulations. This method enables researchers to simulate single-cell transcriptomes before committing to experiments and provides valuable synthetic data for machine learning or deep learning model training applications, potentially accelerating any discovery process.

Related work includes the application of cycleGAN models for transforming gene expression data across modalities. For example, Jeon et al. successfully applied a modified cycleGAN architecture to transform L1000 gene expression profiles into RNA-seq-like profiles^7^. Similarly, Sumeer et al. integrated autoencoders with a cycleGAN architecture to embed data from different modalities into a shared latent space, facilitating cross-modality comparisons^8^. Our work builds on these ideas but focuses on the transformation between bulk RNA-seq and scRNA-seq data, thereby expanding the utility of GAN models in the field of transcriptomics and synthetic data generation. In the sections that follow, we describe our cycleGAN architecture, present training and validation results, and discuss both the advantages and the limitations of this approach—including the ethical implications of using synthetic data, particularly in clinical contexts.

## Results

### 0.1 Model Training and Loss Evaluation

The cycleGAN model was trained for 500 epochs using an 80:20 train:test split on matched bulk and single cell data sets (see Methods for details). Early in training, the generator and discriminator losses declined sharply, followed by a brief fluctuation around epochs 20–25 (Figure 1). The fluctuation observed here reflects the inherent instability in adversarial training, where the model attempts to balance the objectives of the generator and the discriminator^9,10^.

**Figure 1.**
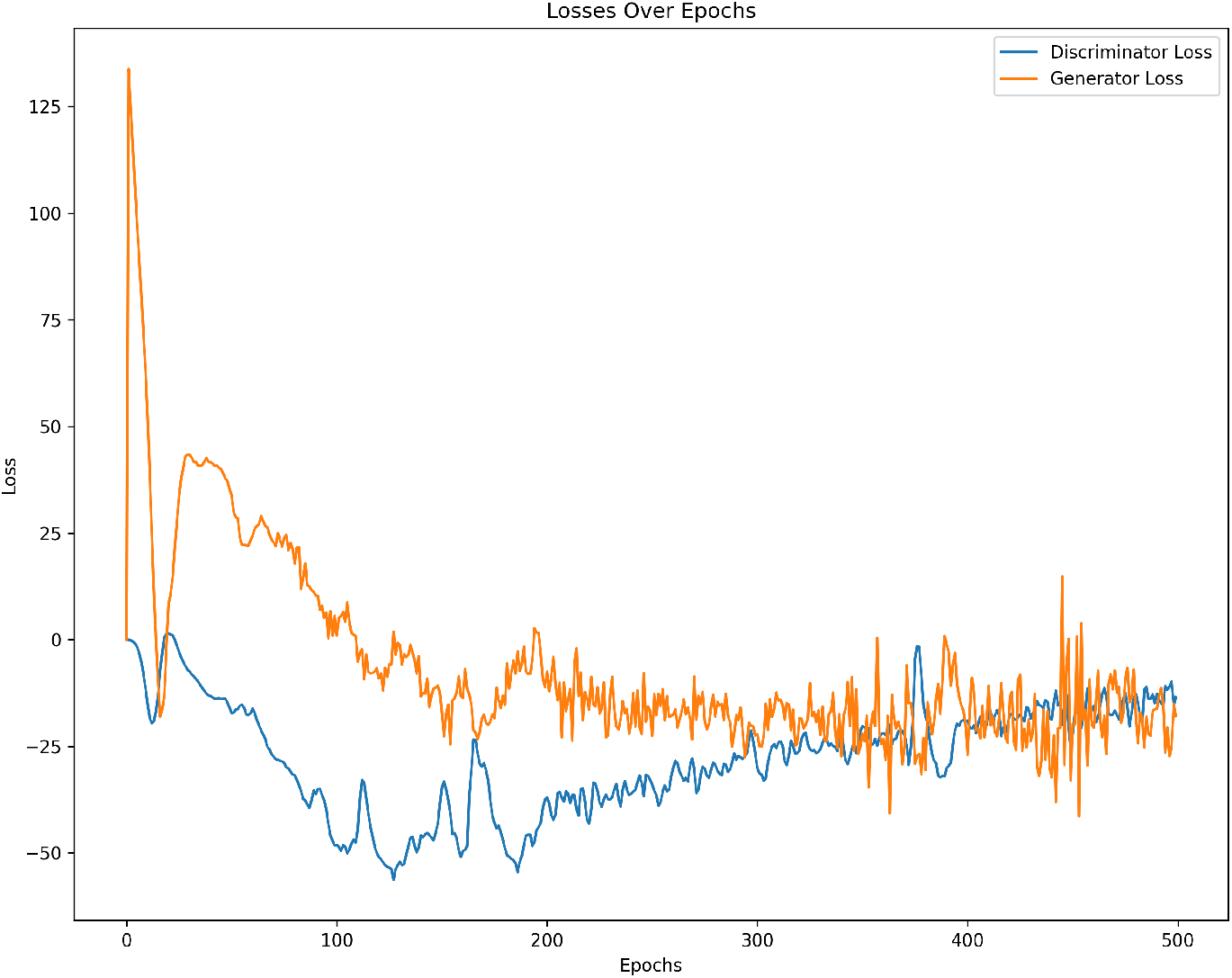
Generator and Discriminator Losses Over 500 Epochs. Following an initial decline, a spike is observed around epochs 20–25, likely reflecting adversarial instability. The training converges after approximately 30 epochs.

After approximately 30 epochs, the adversarial training stabilized and the generator and discriminator losses converged. The final discriminator losses hovered near the values, indicating that real and synthetic data were, on average, difficult to distinguish.

### 0.2 Synthetic scRNA-seq Generation Using Bulk RNA-seq data

To evaluate the trained model, previously unseen bulk RNA-seq profiles were input to generate synthetic single-cell-like data. A Pearson correlation coefficient (PCC) was computed between the generated profiles and the *real* scRNA-seq data from the same training domain. The synthetic data achieved an average PCC of about 0.79 with the real scRNA-seq dataset, suggesting moderate to strong similarity. However, this correlation should not be interpreted as definitive evidence of capturing the *full* cellular heterogeneity.

In Figure 2, we show an example of the generated expression for the NKG7 gene, a marker relevant in certain immune cell populations. Here, the synthetic profiles approximate the real scRNA-seq values, although variability across individual cells remains.

**Figure 2.**
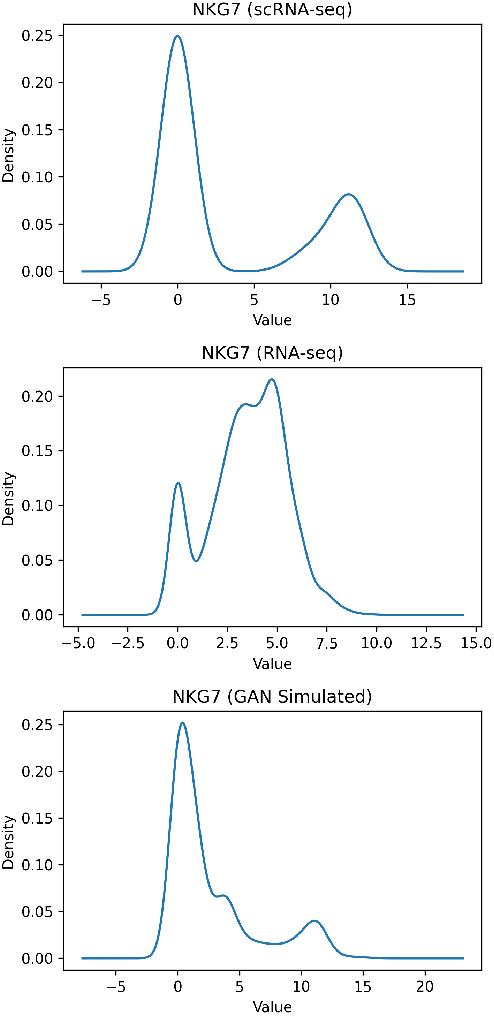
Synthetic Expression Profiles for NKG7. We show a distribution of generated NKG7 expression (blue) compared to real scRNA-seq data (green). The model was trained on T-cell-related datasets in which NKG7 serves as an activation marker.

### 0.3 Principal Component Analysis (PCA)

PCA was performed on real scRNA-seq data, bulk RNA-seq data, and the synthetic scRNA-seq-like profiles to visualize sample-level variability. As shown in Figure 3, the synthetic data (blue) clusters closely to the real single-cell data (green), demonstrating that the model captures some features of the scRNA-seq distribution. However, the synthetic data remain distinct from the original bulk data (red), reinforcing that the cycleGAN performs a nontrivial transformation between the two modalities.

**Figure 3.**
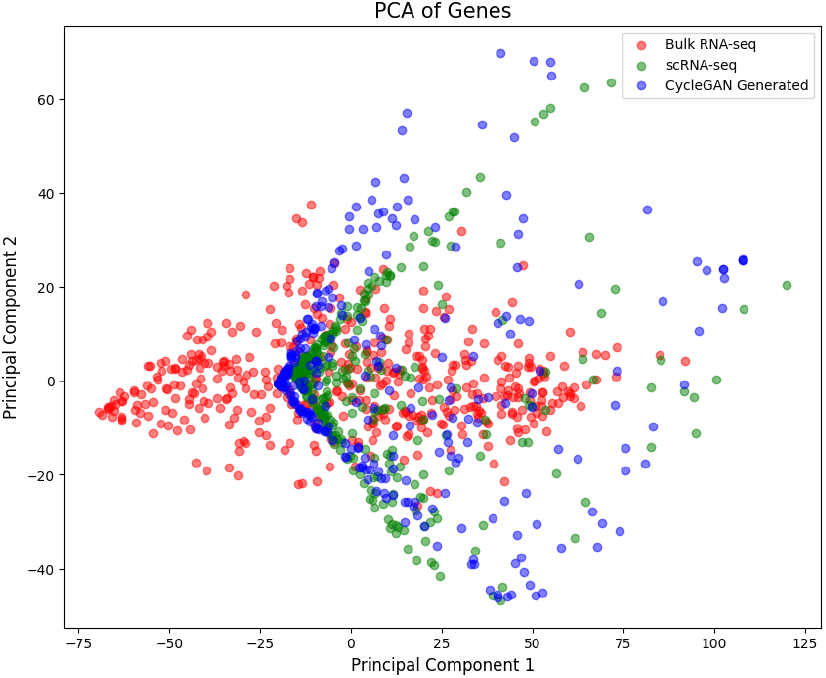
PCA Plot of Real and Synthetic Datasets. The real scRNA-seq data (green) and synthetic single-cell-like data (blue) cluster separately from the bulk RNA-seq data (red), suggesting successful modality transformation.

### 0.4 Performance Metrics

To validate the performance of our model, we calculated the average loss in a held-out test set (400 RNA-seq profiles and 400 scRNA-seq profiles). Table 1 summarizes the average identity loss, cycle consistency loss, and adversarial loss on the held-out test set. Relatively low losses indicate that the transformations preserve domain-specific information and reconstruct the original data reasonably well. However, since we have not tested on completely external or biologically distinct datasets, these metrics primarily reflect the model’s capacity to reconstruct the trained domain rather than to confirm broader generalizability.

**Table 1.** Average losses on the held-out test set. Loss values reflect the model’s reconstruction ability and consistency in domain translation.

| Type | Average Loss |
| --- | --- |
| Average Identity Loss A (Bulk→Bulk) | 0.809 |
| Average Identity Loss B (sc→sc) | 0.779 |
| Average Cycle Loss (A→B→A) | 0.773 |
| Average Cycle Loss (B→A→B) | 0.758 |

### 0.5 Synthetic scRNA-seq Generation Using scRNA-seq Data and Evaluation

To validate the generation of syntehtic transcriptomes, we downloaded bulk RNA-seq profiles of naive CD4+ T immune cells from the DICE data base [16]. The goal for evaluation was to transform the bulk RNA-seq profiles into synthetic single cell RNA-seq profiles and then compare the distributions of the synthetic profiles to naive CD4+ T cells from the single cell study. To further support this comparison, we extracted naive CD4T cell profiles from another scRNA-seq study in order to evaluate them against ghe generate scRNA-seq profiles.

We calculated the dropout rates to evaluate each method’s ability to simulate artificial dropouts. In single-cell RNA-seq data, dropouts refer to instances where a gene’s expression is not detected despite being biologically expressed—primarily due to technical limitations such as low RNA capture efficiency and amplification biases. Synthetic data generation tools aim to replicate this phenomenon by explicitly modeling dropout patterns, injecting “missing” or zero values to mimic both biological variability and technical noise observed in real data. This enables the benchmarking and validation of computational methods under realistic dropout conditions.

Here, we benchmark our GAN-based method against Splatter, a commonly used synthetic data generation tool, as well as a real scRNA-seq dataset. Table 2 presents the dropout rates across all three conditions. While Splatter effectively reproduces overall dropout rates, it fails to capture the underlying gene expression distributions observed in the real dataset.

**Table 2.** Dropout rates for the different methods.

| Method | Dropout |
| --- | --- |
| Splatter | 0.885 |
| Synthetic | 0.793 |
| real | 0.883 |

In addition to evaluating the dropout rates, we further assessed the quality of the synthetic profiles by comparing the mean-variance relationship of the original scRNA-seq dataset with those of the synthetic profiles generated by the GAN-based architecture (labeled as *Synthetic*), Splatter, and ZINB-WaVE. Figure 4 illustrates that the synthetic profiles generated by our GAN architecture closely replicate the mean-variance curve of the real dataset while also capturing gene-specific distributions.

**Figure 4.**
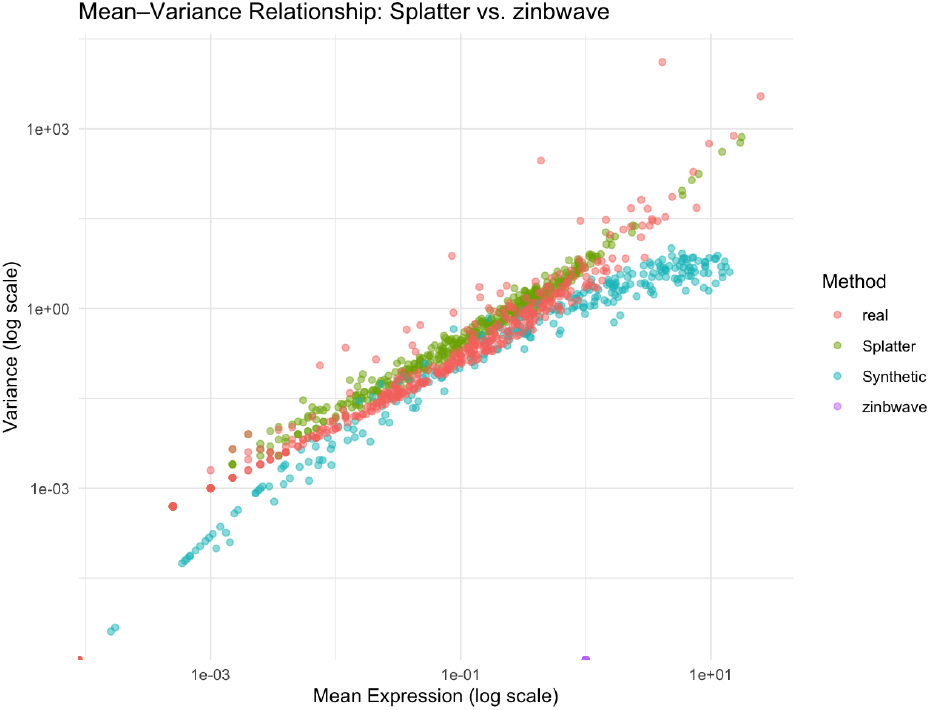
Mean variance plot displaying the mean and variance of the real and synthetic features generated by the models.

From the following evaluations, we can conclude that our method performs on par with pre-available models with the ability to replicate dropout rates from the original dataset as well as capture mean-variance relationships. Additionally, our model also provides the benefit of generating synthetic bulk-RNA-seq data.

## Discussion

In this study, we demonstrate a cycleGAN-based framework to generate synthetic single cellA sequencing (scRNA-seq) data from bulk RNA sequencing (RNA-seq) profiles. This approach addresses a gap in next-generation sequencing by providing a cost-effective tool to generate synthetic single-cell-like data for preliminary investigations and machine learning applications. By learning data distributions and transforming between bulk RNA-seq and scRNA-seq modalities, the cycleGAN achieves a balanced adversarial training process, which enables the generation of synthetic profiles that closely resemble real single-cell data.

The results indicate that the synthetic data generated capture many features represented in the scRNA-seq data, including their characteristics such as sparsity and variation. The gene expression density plots and PCA show that the synthetic profiles match the distributions of the real single-cell profiles. This highlights the potential utility of this method for cost-effective preliminary exploratory studies. Despite a relatively strong Pearson correlation of approximately 0.79, further improvements could include incorporating more diverse training datasets to better capture variations across different cell types and biological conditions, as well as experimenting with other loss functions and regularization strategies (e.g., gradient penalty or spectral normalization) to mitigate common instability issues in GAN-based models.

To further evaluate the synthetic profiles generated by our GAN based model, we bench-marked it against one other publicly available synthetic scRNA-seq data generation tools. We evaluated the synthetic profiles by measuring how well they were able to capture dropouts and by looking at their mean-variance statistics. Our GAN based model generated synthetic profiles with 79 percent droupout, close to the ideal rates observed in a scRNa-seq dataset. Moreover, our model was able to effectively capture the distributions of the gene expression compared to the synthetic profiles generated by Splatter which does not represent expression levels in a gene-specific manner.

In addition to providing a method for simulating synthetic cell-level transcriptomes, our framework can help researchers conduct in silico experiments such as injecting synthetic profiles as features to address class imbalances or serve as augmentation in deep learning training protocols. Importantly, however, the synthetic data cannot substitute for true single-cell experiments. Bulk RNA-seq lacks the resolution and precise cell-type proportions of rare subpopulations, and any variation not present in the training dataset will likely be misrepresented in the synthetic output. Consequently, the method should be viewed as a tool to supplement data augmentation techniques and not as a technique that would replace experimental scRNA-seq.

While synthetic data can address privacy concerns by abstracting patient-specific details, researchers must exercise caution about their use in clinical contexts, where the nuances of real patient heterogeneity and disease state require careful validation. Misinterpretation of synthetic profiles could lead to inaccurate conclusions. More benchmarking against external datasets will help established and define the strengths, limitations, and reproducibility of the method in various biological settings. Batch effects between bulk RNA-seq and scRNA-seq platforms may influence model training and we have tried to minimize batch effects by using matched biological contexts (same cell types/conditions).

The scalability of this framework is another important consideration. Although the approach successfully handles moderate dataset sizes, larger and more complex transcriptomic datasets require additional computational resources, such as more powerful GPUs and efficient data loading pipelines.

## Conclusions

Our cycleGAN-based approach offers a cost-effective way to generate scRNA-seq-like transcriptomic profiles from bulk RNA-seq data, allowing for preliminary single cell exploratory analyzes and data augmentation for deep learning tasks. However, this method does not replace real scRNA-seq or bulk RNA-seq experiments because synthetic profiles do not capture all aspects of cellular diversity. The model’s ability to generate unseen cell types or new transcriptional states is inherently limited by the scope of its training data, and multiple synthetic solutions may arise from the same bulk sample. Additionally, new benchmarking methodologies need to be established in order to determine the broader validity of the tool. Future work will focus on expanding the training set, improving model stability and scalability, and carefully evaluating the ethical implications of using synthetic data, especially in clinical contexts. By addressing these challenges, our framework has the potential to serve as a practical tool for exploring single-cell heterogeneity and advancing transcriptomic research in a variety of biological and medical applications.

## Methods

### 0.6 Data Acquisition and Preprocessing

The datasets were sourced from the ARCHS4 RNA-seq repository (bulk)^12^ and a publicly available scRNA-seq study investigating cardiovascular disease^15^. We curated a total of 2,000 RNA-seq profiles and 2,000 single-cell profiles representing the same biological domain (e.g., CD4+ T cell subsets). After filtering for genes (461 genes) profiled across both modalities, we identified 461 overlapping genes.

An 80:20 train:test split (400 test profiles) was then applied within each modality, ensuring that the test set remained unseen during training. Data were normalized by counts per million (CPM) and subsequently log-transformed to stabilize variance across genes.

### 0.7 CycleGAN Architecture and Training

We adopt the standard cycleGAN framework^10^, comprising two generators (*G*_*A*→*B*_, *G*_*B*→*A*_) and two discriminators (*D*_*A*_, *D*_*B*_). Figure 5 shows a schematic diagram of the model’s training procedure:

**Figure 5.**
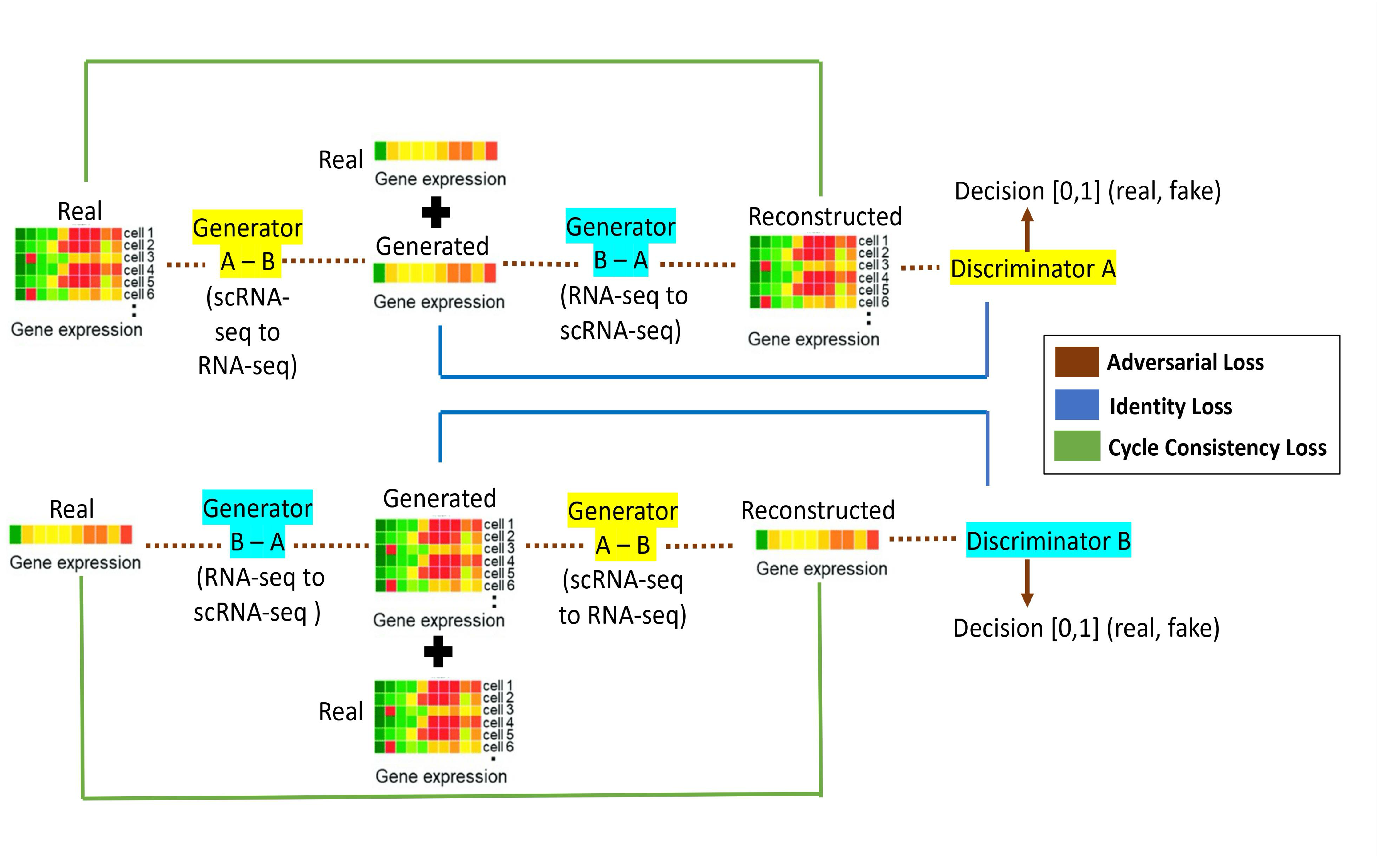
Schematic diagram of the network’s training procedure.

- *G*_*A*→*B*_: Translates bulk RNA-seq (Domain A) to synthetic scRNA-seq (Domain B).
- *G*_*B*→*A*_: Translates scRNA-seq (Domain B) back to bulk RNA-seq (Domain A).
- *D*_*A*_: Discriminates between real and synthetic bulk RNA-seq.
- *D*_*B*_: Discriminates between real and synthetic single-cell RNA-seq.

### Model Architecture and Training

#### 0.8.1 Optimization and Learning Setup

The learning rate plays a critical role in gradient-based optimization, directly impacting model convergence. For our CycleGAN framework, we set the learning rate to 5 × 10^−5^ for the generators and 1 × 10^−5^ for the discriminators. This conservative choice ensures stable weight updates, preventing rapid oscillations or divergence during training. To further stabilize training dynamics, we employed the Adam optimizer with beta coefficients of (*β*_1_, *β*_2_) = (0.5, 0.999). This parameterization balances model responsiveness to gradient updates while preserving long-term memory of past gradients, facilitating smooth and effective convergence.

#### 0.8.2 Regularization and Loss Functions

To balance the adversarial objectives of identity loss and cycle consistency loss, we assigned loss weighting coefficients of and 25, respectively. The greater emphasis on cycle consistency loss underscores the importance of preserving biological context to a certain extent when transforming between RNA-seq and scRNA-seq modalities. Additionally, we imposed weight clipping constraints in the range [− 0.01, 0.01] on the discriminator to avoid exploding gradients under the Wasserstein loss function. These regularization strategies ensure robust adversarial training and mitigate mode collapse.

Several loss functions were used to drive the model training process, Total Generator Loss: The total loss for the generator is a weighted sum of the identity loss, adversarial losses, and cycle consistency losses.

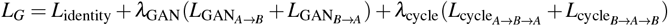

1. The **identity loss** was calculated to verify that data passed through the generator of the same domain remains consistent.

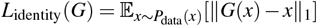

2. The **adversarial Loss** is used to drive generators to produce data that is indistinguishable from real images by the discriminators.

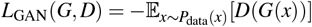

3. The **cycle consistency loss** was measured to ensure that transformations between RNA-seq and scRNA-seq modalities retained the original data structure when converted back. Additionally, Pearson correlation coefficients were computed to assess the similarity between the generated scRNA-seq profiles and real data. Mean absolute error (L1 norm) and Wasserstein loss were used to track the performance of both the generators and discriminators over time.

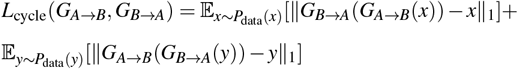

Total **adversarial loss for discriminators** is used to drive discrimination between real and fake data. The loss is the difference between the discriminator’s predictions on real and fake data, and consists of two mirrored terms for discriminators A and B respectively.

For discriminator A:

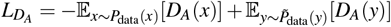

For discriminator B:

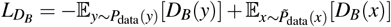

Each of these losses is back-propagated through the network to update the model parameters during the training process.

#### 0.8.3 Network Design and Training Protocol

The generator was constructed using a modified ResNet50 architecture, leveraging residual blocks to enhance representational learning. The encoder mapped input gene expression profiles into a latent space, while the decoder reconstructed the transformed output. The discriminator was implemented as a deep feed-forward network with ReLU activations that evaluated the realism of generated samples. The adversarial interplay between these components refined the synthetic data quality, promoting biologically plausible mappings between modalities.

The model was trained for 500 epochs with a batch size of 200 samples per epoch. Training was conducted using the Adam optimizer, with mini-batching to optimize computational efficiency and convergence. The dataset, comprising paired scRNA-seq and RNA-seq profiles, was evaluated on a held-out test set to evaluate model performance and generalization. These methodological choices collectively ensure stable training and synthetic data generation.

### Hyperparameter Settings

- **Learning Rate:** 0.00005 (generators), 0.00001 (discriminators).
- **Adam Optimizer Betas:** (0.5, 0.999).
- **Loss Weights:** *λ*_GAN_ = 1, *λ*_cycle_ = 25, identity loss scaling = 0.5.
- **Batch Size:** 200 samples/epoch, for a total of 500 epochs.

The model was implemented in PyTorch^14^ and trained on an NVIDIA V100 GPU (32GB RAM). Training took approximately 3 hours.

## Author contributions statement

Conceptualization, S.S.A.; methodology, S.S.A.; software, S.S.A.; validation, S.S.A.; formal analysis, S.S.A.; investigation, D.R.; data curation, S.S.A.; writing—original draft preparation, S.S.A.; writing—review and editing, S.S.A. and D.R.; visualization, S.S.A.; supervision, D.R.; project administration, D.R.; funding acquisition, D.R. All authors have read and agreed to the published version of the manuscript.

## Additional information

### Data Availability

All relevant code and data used to train and evaluate this model are publicly available on GitHub (https://github.com/sujitsilas/synthetic-transcriptome-generator/). The bulk RNA-seq data were obtained from the ARCHS4 repository (https://maayanlab.cloud/archs4/index.html), while the scRNA-seq data were sourced from GEO (accession number: GSE190570)^15^. For additional data or information, please contact Sujit Silas Armstrong.

